# Post-transcriptionally impaired *de novo* mutations contribute to the genetic etiology of four neuropsychiatric disorders

**DOI:** 10.1101/175844

**Authors:** Fengbiao Mao, Lu Wang, Xiaolu Zhao, Zhongshan Li, Luoyuan Xiao, Rajesh C. Rao, Jinchen Li, Huajing Teng, Xin He, Zhong Sheng Sun

## Abstract

While deleterious *de novo* mutations (DNMs) in coding region conferring risk in neuropsychiatric disorders have been revealed by next-generation sequencing, the role of DNMs involved in post-transcriptional regulation in pathogenesis of these disorders remains to be elucidated. Here, we identified 1,736 post-transcriptionally impaired DNMs (piDNMs), and prioritized 1,482 candidate genes in four neuropsychiatric disorders from 7,748 families. Our results revealed higher prevalence of piDNMs in the probands than in controls (*P* = 8.19×10^−17^), and piDNM-harboring genes were enriched for epigenetic modifications and neuronal or synaptic functions. Moreover, we identified 86 piDNM-containing genes forming convergent co-expression modules and intensive protein-protein interactions in at least two neuropsychiatric disorders. These cross-disorder genes carrying piDNMs could form interaction network centered on RNA binding proteins, suggesting a shared post-transcriptional etiology underlying these disorders. Our findings illustrate the significant contribution of piDNMs to four neuropsychiatric disorders, and lay emphasis on combining functional and network-based evidences to identify regulatory causes of genetic disorders.

## Introduction

Next-generation sequencing, which allows genome-wide detection of rare and de novo mutations (DNMs), is transforming the pace of genetics of human disease by identifying protein-coding mutations that confer risk^1^. Various computational methods have been developed to predict the effects of amino acid substitutions on protein function, and to classify corresponding mutations as deleterious or benign, based on evolutionary conservation or protein structural constraints^2, 3^. Beside the effect on protein structure and function, genetic mutations involve in transcriptional processes^4, 5^ via direct or indirect effects on histone modifications^6^, and enhancers^7^ to affect pathogenesis of diseases. However, the majorities of mutation are located in non-coding regions, and some of them have no relationship with transcriptional regulation, but can lead to an observable phenotype or disease^8^, suggesting the existence of another layer of regulatory effect of mutations. It has been revealed that single nucleotide variants can alter RNA structure, known as RiboSNitches, and depletion of RiboSNitches result in the alteration of specific RNA shapes at thousands of sites, including 3’untranslated region, binding sites of RBPs and microRNAs^9^. Thus, the mutations can impair post-transcriptional processes through disrupting the binding of micRNAs and RNA binding proteins (RBPs)^10, 11^, resulting in various human diseases. For example, a variant in the 3’ untranslated region of FMR1 decreases neuronal activity-dependent translation of FMRP by disrupting the binding of HuR, leading to developmental delay in patients^10^. Some attempts have been undertaken to better understand the interactions between mutations and binding of noncoding RNAs or RBPs. Maticzka *et al*. developed a machine learning-based approach to predict protein binding sites on RNA from crosslinking immunoprecipitation (CLIP) data using both RNA structure and sequence features^12^. Fukunaga *et al*. developed the CapR algorithm based on the probability of secondary structure of an RNA for RBP binding^13^. However, identifying the network between single nucleotide mutations and post-transcriptional regulation remains challenging because of the complexity of the underlying interaction networks. Our and other’s methods named RBP-Var^14^ and POSTAR^15^ represent initial efforts to systematically annotate post-transcriptional regulatory maps, which hold great promise for exploring the effect of single nucleotide mutations on post-transcriptional regulation in human diseases.

Increasing prevalence of neuropsychiatric disorders in children with unclear etiology has been reported during the past three decades^16^. Whole-exome sequencing of pediatric neuropsychiatric disorders uncovered the critical role of DNMs in the pathogenesis of these disorders^1^. However, previous studies of these disorders have focused on mutations in coding region^1^, cis-regulation^17, 18^, epigenome^19^, transcriptome^20, 21^, and proteome^22^, very few is known about the effect of DNMs on post-transcriptional regulation. Recently, more attentions have been paid on DNMs in regulatory elements and non-coding regions in neurodevelopmental disorders as it is indispensable to combine functional and evolutionary evidence to identify regulatory causes of genetic disorders^23, 24^. Most recently, a deep-learning-based framework illuminates involvement of noncoding DNMs in synaptic transmission and neuronal development in autism spectrum disorder^25^. Therefore, it is imperative to identify the post-transcriptionally regulation-disrupting DNMs related to pathology and clinical treatment of neuropsychiatric disorders.

To test whether post-transcriptionally regulation-disrupting DNMs contribute to the genetic architecture of psychiatric disorders, we collected whole exome sequencing data from 7,748 core families (5,677 families were parent-probands trios and 2,071 families were normal trios) and curated 9,519 *de novo* mutations (6,996 DNMs in probands and 2,523 DNMs in controls) from four kinds of neuropsychiatric disorders, including autism spectrum disorder (ASD), epileptic encephalopathy (EE), intellectual disability (ID), schizophrenia (SCZ), as well as unaffected control subjects (Supplementary Table 1). By employing our newly updated workflow RBP-Var2 (Figure 1A, Supplementary Table 2) from our previously developed RBP-Var^14^, we investigated the potential impact of these *de novo* mutations involved in post-transcriptional regulation in these four neuropsychiatric disorders based on experimental data of genome-wide association studies (GWAS), expression quantitative trait locus (eQTL), CLIP-seq derived RBP binding sites, RNA editing and miRNA targets, and found that a subset of *de novo* mutations could be classified as post-transcriptionally impaired DNMs (piDNMs). These piDNMs showed significant enrichment in cases after correcting for multiple testing, and genes hit by these piDNMs were further analyzed for their properties and relative contribution to the etiology of neuropsychiatric disorders.

**Figure 1.**
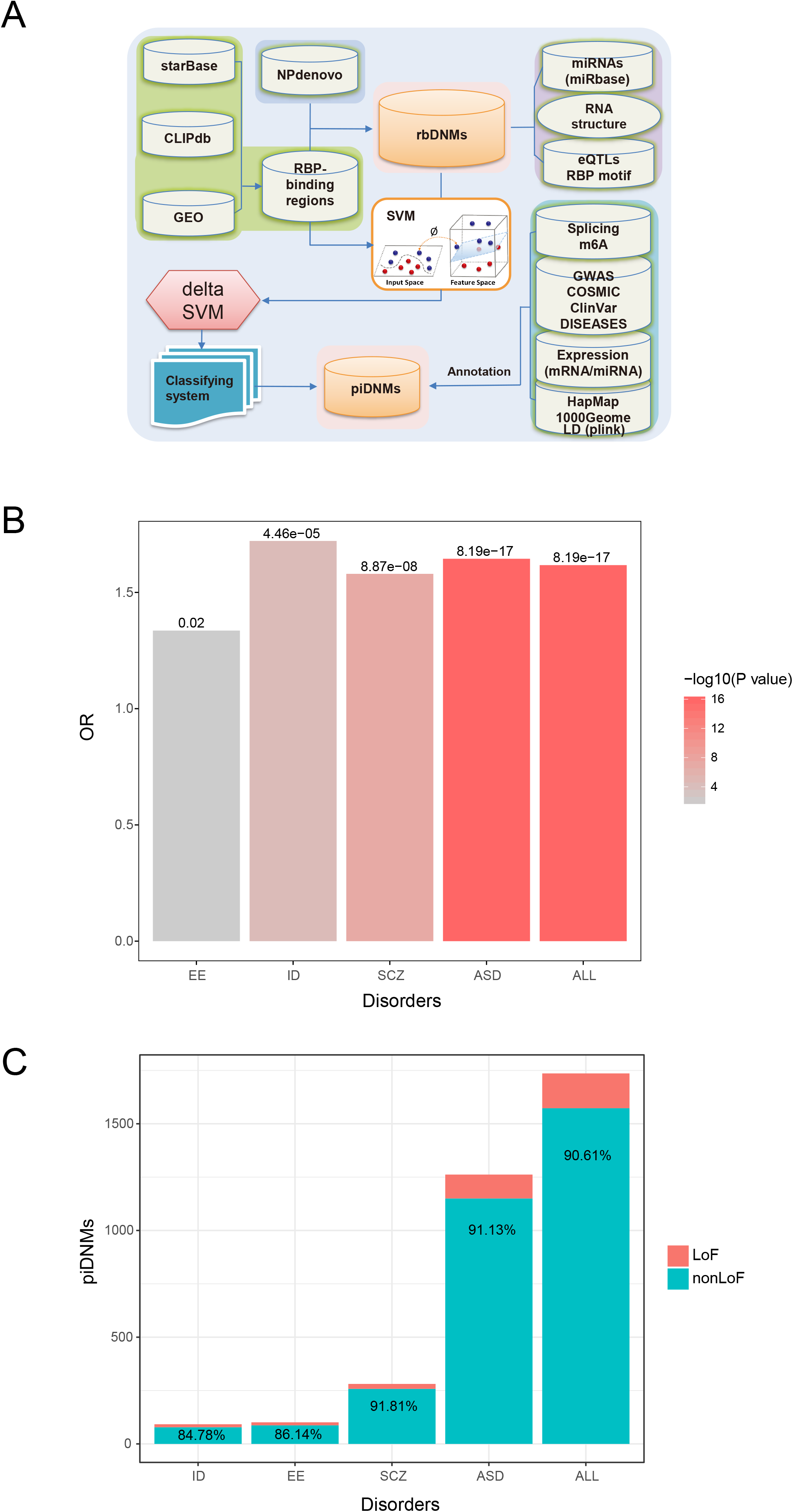
The abundance of piDNMs in different disease categories. (A) Schematic overview of the RBP-Var2 algorithm. (B) The bar plot corresponds to the odds ratios indicating the enrichment of piDNMs in patients from each of the four neuropsychiatric disorders. (C) The relative amount of LoF and non-LoF piDNMs in five neuropsychiatric disorders.

## Results

### The frequency of piDNMs is much higher in probands than that in controls

To test whether specific subsets of regulatory DNMs contribute to the genetic architecture of neuropsychiatric disorders, we devised and updated the method, RBP-Var2 (http://www.rbp-var.biols.ac.cn/), based on experimental data of GWAS, eQTL, CLIP-seq derived RBP binding sites, RNA editing and miRNA targets. Subsequently, we used our updated workflow to identify functional piDNMs from 5,677 trios with 6,996 DNMs across four neuropsychiatric disorders as well as 2,071 unaffected controls with 2,523 DNMs (Supplementary Table 1). We determined DNMs with 1/2 category score predicted by RBP-Var2 as piDNMs when considering their impact on RNA secondary structure, the binding of miRNAs and RBPs, and identified 1,736 piDNMs in probands (Supplementary Table 3), of which 17,7,7,6 and 1,699 were located in 3’ UTRs, 5’ UTRs, ncRNA exons, splicing sites and exons, respectively. In detail, RBP-Var2 identified 1,262 piDNMs in ASD, 281 piDNMs in SCZ, 101 piDNMs in EE, 92 piDNMs in ID and 354 piDNMs in healthy controls (Supplementary Table 3, 4). Interestingly, the frequency of piDNMs in the four neuropsychiatric disorders were significantly over-represented compared with those in controls (OR = 1.62, *P* = 8.19×10^−17^, Table 1). We also observed that probands groups have much more abundant piDNMs compared with controls in four kinds of neuropsychiatric disorders (Figure 1B; Table 1). Dramatically, we found that synonymous piDNMs were significantly enriched in probands in contrast to those in controls (*P*=9.73×10^−4^). While in the data set of original DNMs before the evaluated by RBP-Var2, the enrichment of the synonymous DNMs was not observed in cases, which is consistent with previous study^1^. To eliminate the effects of loss-of-function (LoF) mutations, we filtered out those LoF mutations from all piDNMs and found the non-LoF piDNMs also exhibited higher frequency in probands (*P*=2.36×10^−14^) (Table 1, Figure 1C; Supplementary Figure 1). Our analysis found a subset of DNMs, namely piDNMs, are enriched in probands and may contribute to the pathogenesis of these disorders although the rate of all de novo synonymous variants, which as a category, does not contribute significantly to risk for neurodevelopmental disorders.

### The piDNMs outperforms protein-disruptive DNMs in risk prediction

To investigate the accuracy and specificity of DNMs in different regulatory processes, we compared our tool with other three variant effect prediction tools, including SIFT^25^, PolyPhen2 (PPH2)^26^ and RegulomeDB^26^. We found that the frequency of the stop gain DNMs is higher in cases than in controls determined by SIFT (*P =* 4.83×10^−2^), and higher frequency of nonsynonymous DNMs was identified by PPH2 (*P* = 1.82×10^−2^) (Figure 2A, B). However, RegulomeDB determined no significant higher frequency of functional DNMs in any functional category (Figure 2C) in cases versus controls. In contrast, RBP-Var2 could determine much more functional DNMs in the categories of frameshift (*P =* 1.38×10^−3^), nonsynonymous (*P =* 8.79×10^−15^), stopgain (*P =* 6.42×10^−4^) and synonymous (*P =* 7.30×10^−4^) (Figure 2D). Then, we performed receiver operating characteristic (ROC) analysis to systemically evaluate the sensitivity and specificity of these four prediction methods. We found that area under curve (AUC) value of SIFT, PPH2, RBP-Var2 and RegulomeDB are 78.27%, 76.57%, 82.89% and 50.77%, respectively (Supplementary Figure 2), indicating that SIFT, PPH2, and RBP-Var2 is more sensitive and specific than that of RegulomeDB with *P* value 1.63×10^−10^, 2.40×10^−8^ and 2.51×10^−60^, respectively. In addition, the AUC value of RBP-Var2 is higher than that of SIFT and PPH2 with *P* value 0.049 and 0.019, respectively. Intriguingly, RBP-Var2 could detect an additional 928 piDNMs covering 665 genes that were regarded as benign DNMs by other three methods, accounting for 25.27% of total 3,672 deleterious DNMs detected by all four tools (Supplementary Figure 3A, B). Especially, the non-LoF piDNMs detected by RBP-Var2 alone account for 52.8% of non-LoF piDNMs, while only 26.2% of non-LoF piDNMs were classified to be deleterious predicted by both SIFT and Polyphen2 (Supplementary Figure 3C). The top three enriched gene ontology of these 665 genes were intracellular signal transduction (*P* = 7.41×10^−6^), organelle organization (*P* = 8.90×10^−6^) and mitotic cell cycle (*P* = 2.06×10^−5^) (Supplementary Figure 3D; Supplementary Table 5), suggesting dysregulation involved in cell cycle and the impaired signal transduction may contribute to diverse neural damage, thereby trigger neurodevelopmental disorders^27, 28^. Therefore, the piDNMs detected by RBP-Var2 are distinct, and may play significant roles in the post-transcriptional processes of the development of neuropsychiatric disorders.

**Figure 2.**
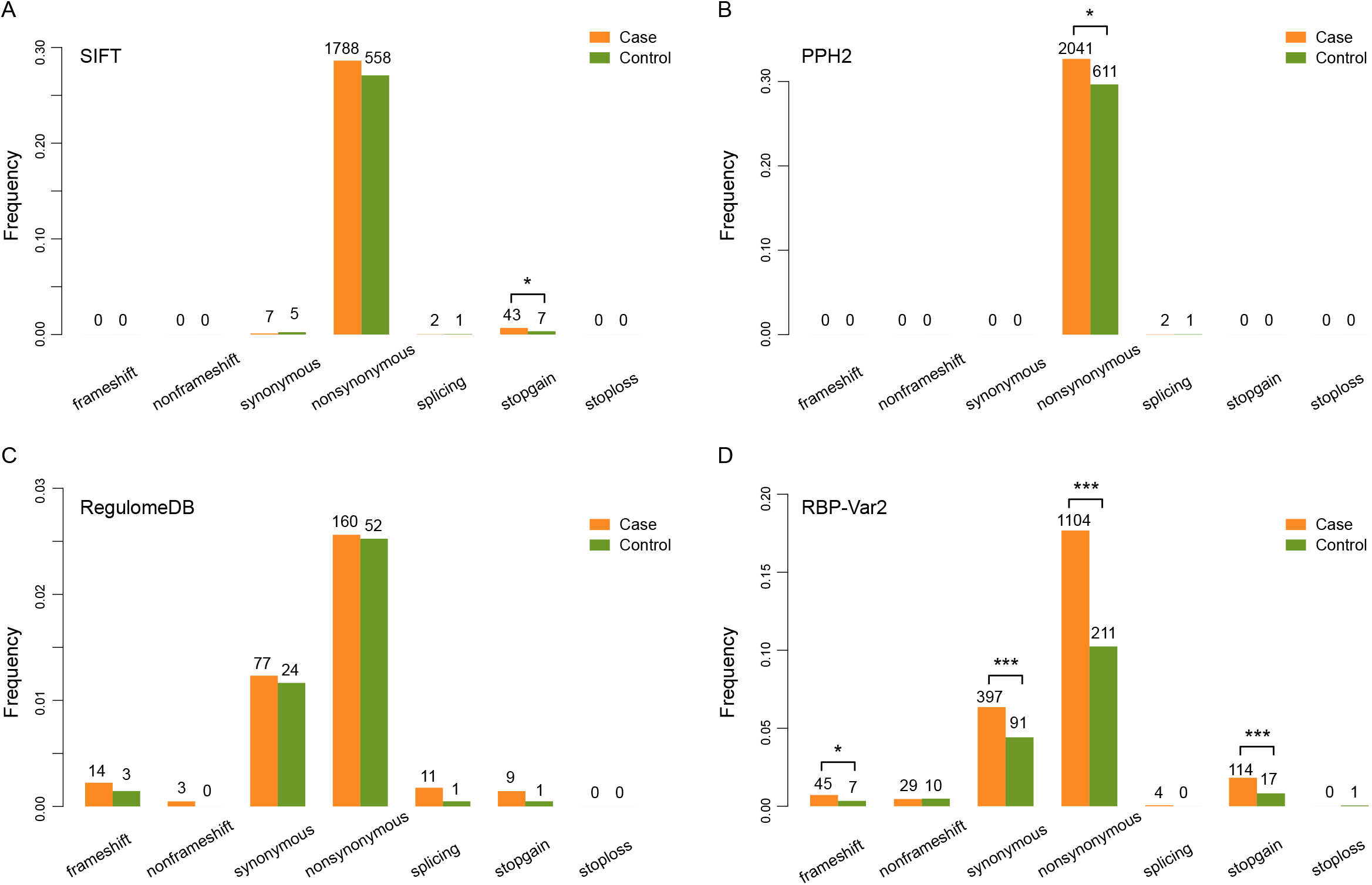
Performance comparison of the ability to distinguish severe DNMs between RBP-Var2 and three other tools. (A) Different kinds of DNMs affecting protein function predicted by SIFT. The Y-axis corresponds to the proportion of each kind of mutations within the total number of damaging DNMs predicted by SIFT. (B) Different kinds of DNMs that affect protein function predicted by PolyPhen2. The Y-axis corresponds to the proportion of each kind of mutations within the total number of damaging DNMs predicted by PolyPhen2. (C) The DNMs, predicted as functional elements involved in transcriptional regulation by RegulomeDB, are categorized into different functional types. The Y-axis corresponds to the proportion of each kind of mutations within the total number of damaging DNMs predicted by RegulomeDB. (D) The DNMs classified as either level 1 or 2 (piDNMs) are categorized into different functional types. The Y-axis corresponds to the proportion of each kind of mutations within the total number of damaging piDNMs. The *P* values were measured by two-sided binomial test. DNMs predicted in both cases and controls are excluded in the comparison and the DNMs labeled as “unknown” are not demonstrated in the bar plot.

### Genes hit by piDNMs are shared across four neuropsychiatric disorders

Firslty, we identified 13 recurrent piDNMs, including seven piDNMs in ASD and six piDNMs in ID (Figure 3A). Secondly, we identified 149 genes carrying at least two piDNMs in all disorders, including 128 genes in ASD, three genes in EE, ten genes in ID and eight in SCZ. Among these 149 genes, we identified 21 high risk genes with *P* value < 1×10^−2^ derived from our previously published TADA program (Transmission And De novo Association)^29^ (Figure 3B). As our previous study using the NPdenovo database demonstrated that DNMs predicted as deleterious in the protein level are shared by four neuropsychiatric disorders^30^, we then wondered whether there were common piDNMs among four neuropsychiatric disorders. By comparing the genes harboring piDNMs across four disorders, we found 86 genes significantly shared by at least two disorders rather than random overlaps (permutation test, *P <*1.00×10^−5^ based on random resampling, Figure 3C, D). Similar results have been observed for the overlap between the cross-disorder genes of any two/three disorders and the genes in control, as well as for the overlapping genes between each disorder and the control (Supplementary Figure 4B-O). In addition, the numbers of shared genes for any pairwise comparison or any three disorders are all significantly higher than randomly expected except for the comparison of EE versus SCZ (*P=*0.4964) (Supplementary Figure5). Our observation revealed the existence of common genes harboring piDNMs among these four neuropsychiatric disorders.

**Figure 3.**
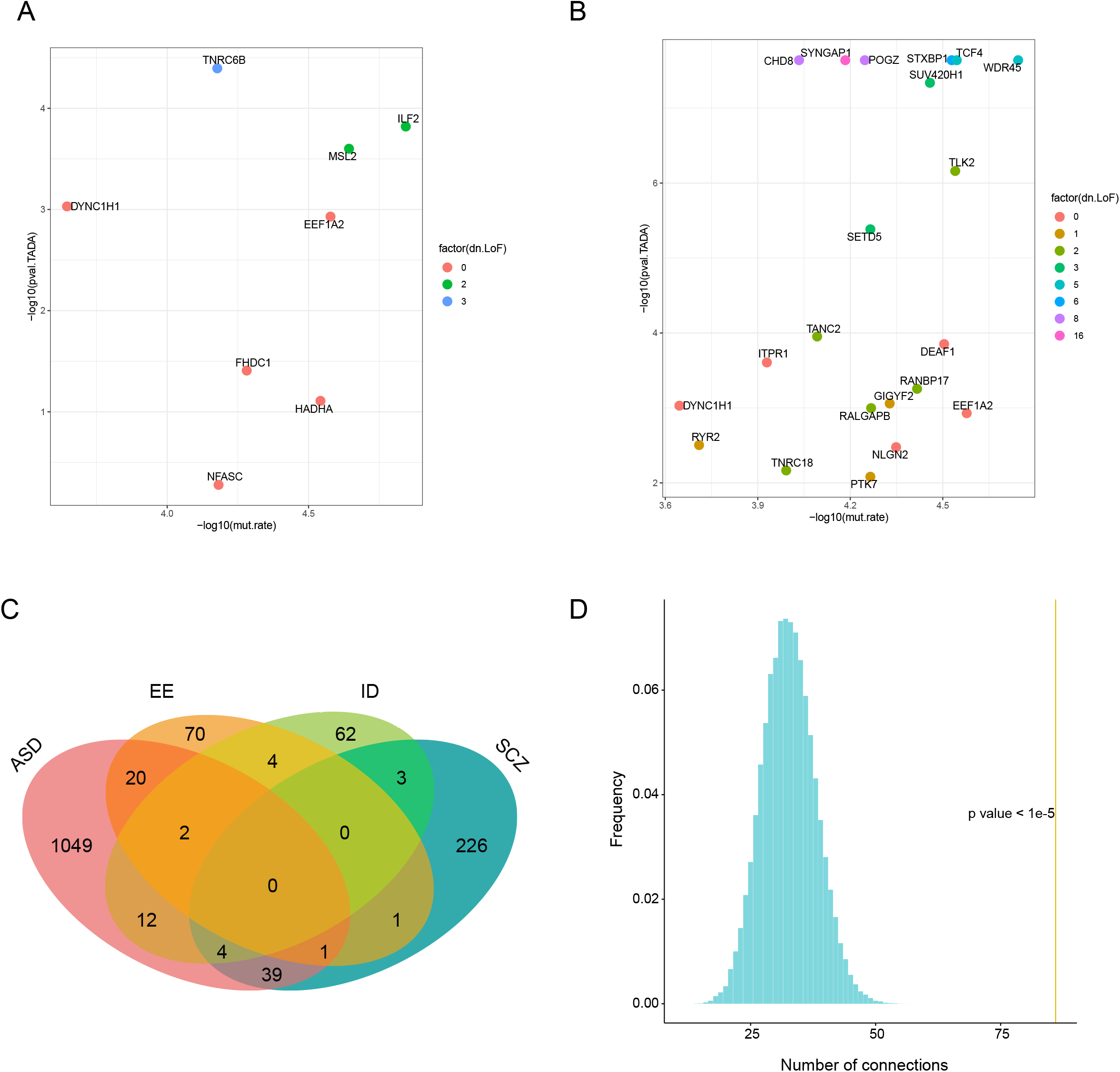
Genes with piDNMs involved in four neuropsychiatric disorders. (A) Scatter plot of eight genes harboring recurrent piDNMs among 1,736 piDNMs. The Y-axis corresponds to the -log_10_(*P* value) calculated by TADA. The X-axis stands for the TADA output of -log_10_(mutation rate). (B) Scatter plot of 21 recurrent genes with p < 0.01 by TADA. The Y-axis corresponds to the -log_10_(*P* value) calculated by TADA. The X-axis stands for the TADA output of -log_10_(mutation rate). (C) Venn diagram representing the distribution of candidate genes shared among the four neuropsychiatric disorders. (D) Permutation test for the randomness of the overlap between the 86 cross-disorder genes. We shuffled the genes of each disorder and calculated the shared genes between the four disorders, and repeated this procedure for 100,000 times to get the null distribution. The vertical dash line indicates the observed value.

### Genes harboring piDNMs are involved in epigenetic modification and synaptic functions

The phenomenon of shared genes among the four neuropsychiatric disorders suggest there may exist common molecular mechanisms underlying their pathogenesis. Thus, we performed functional enrichment analysis for these shared genes, and found they were remarkably enriched in biological processes in chromatin modification like histone methylation, functional classifications of neuromuscular control and protein localization to synapse (Supplementary Table 5, Supplementary Figure 6). These epigenetic regulating genes are composed of CHD5, DOT1L, JARID2, MECP2, PHF19, PRDM4 and TNRC18 (Supplementary Table 6). Moreover, most of these epigenetic modification genes, have been previously linked with neuropsychiatric disorders^31–35^. Interestingly, shared genes in enrichment analyses have intensive linkages among these significant pathways as some of piDNM-containing genes could play roles in more than one of these pathways (Supplementary Figure 7).

Next, to investigate the biological pathways involved in each group of disorder-specific genes with piDNMs, we carried out functional enrichment analysis with terms in biological process (Supplementary Table 7-9). The top three enriched categories of ASD-specific genes were “macromolecule modification” (*P* = 2.90×10^−13^), “organelle organization” (*P* = 1.22×10^−11^) and “cell cycle” (*P* = 1.17×10^−9^). With respect to genes specific to SCZ, it is actually no surprise that the significantly enriched categories are related to protein localization and calcium transport, which have been revealed to be involved in the pathophysiology of schizophrenia^36^. Because of the limited number of genes, only two GO terms were enriched for EE-specific genes, which were “N-glycan processing” (*P* = 3.65×10^−5^) and “protein deglycosylation” (*P* = 2.17×10^−4^), while no terms were statistically enriched for ID-specific genes. Our observation that each group of disorder-specific genes being overrepresented into different biological pathways, suggests that piDNMs may also play a role in the distinct phenotypes of the four psychiatric disorders although the explicit underlying mechanisms need to be further explored.

### Co-expression modules are convergent for cross-disorder genes hit by piDNMs

Co-expression of genes can be used to explore the common and distinct molecular mechanisms in neuropsychiatric disorders^37^. Thus, we performed weighted gene co-expression network analysis (WGCNA)^38^ for the 86 cross-disorder piDNMs-containing genes based on gene expression in 16 human brain structures across 31 developmental stages from BrainSpan developmental transcriptome (n=524)^39^. The results of WGCNA deciphered two gene modules with distinct spatiotemporal expression patterns (Figure 4A, B; Supplementary Figure 8). The turquoise module (n=55 genes) was characterized by high expression during early fetal development (8-24 postconceptional weeks) in the majority of brain structures (Figure 4C). Whereas, the blue module (n=22 genes) showed low expression in early fetal development (8-38 postconceptional weeks) in the majority of brain structures (Figure 4D). It is also crucial to clarify gene expression of these genes in early development stages since altered epigenetic regulation in early development has been shown to be associated with neurodevelopmental disorders^40^. We found most of these 86 genes are highly expressed and may be required for the normal development of human embryo (Figure 4E, Supplementary Table 10). Our observation indicates that these cross-disorder piDNMs-containing genes may play important roles in not only early brain developmental but also early embryonic development.

**Figure 4.**
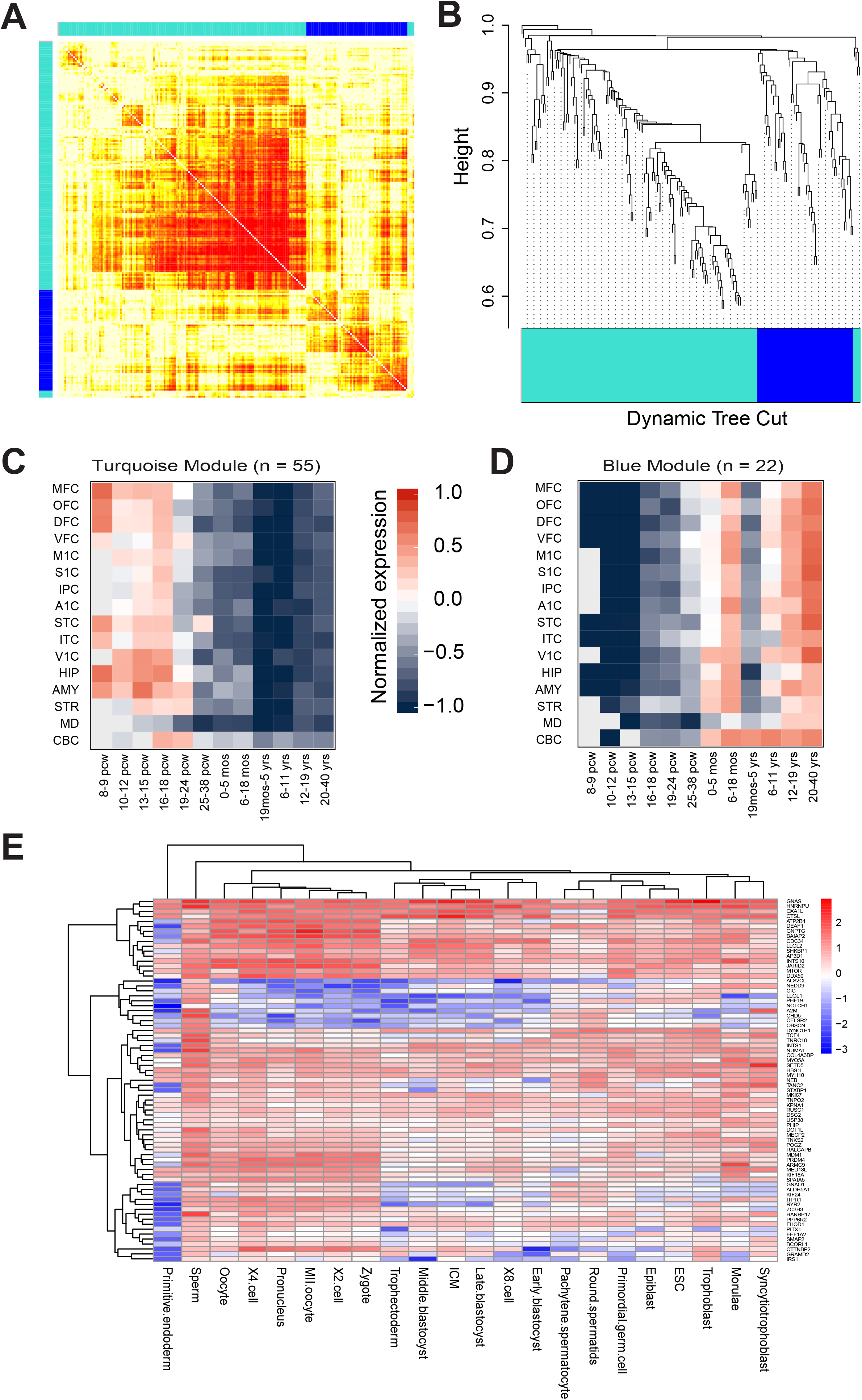
Weighted co-expression analysis of 86 shared genes. (A) Heatmap visualization of the co-expression network of 86 shared genes. The more saturated color corresponds to the more highly expressed genes. (B) Hierarchical clustering dendrogram of the two color-coded gene modules displayed in (A). (C, D) The two spatiotemporal expression patterns (Turquoise module and Blue module) for network genes based on RNA-seq data from BrainSpan, and they correspond to 17 developmental stages across 16 subregions: A1C, primary auditory cortex; AMY, amygdaloid complex; CBC, cerebellar cortex; DFC, dorsolateral prefrontal cortex; HIP, hippocampus; IPC, posteroinferior parietal cortex; ITC, inferolateral temporal cortex; M1C, primary motor cortex; MD, mediodorsal nucleus of thalamus; MFC, anterior cingulate cortex; OFC, orbital frontal cortex; STC, posterior superior temporal cortex; STR, striatum; S1C, primary somatosensory cortex; V1C, primary visual cortex; VFC, ventrolateral prefrontal cortex.

### Protein-protein interactions are intensive for cross-disorder piDNM-containing proteins

The co-expression results indicate that the proteins coded by the 86 cross-disorder genes may have intensive protein-protein interactions (PPIs). To identify common biological processes that potentially contribute to disease pathogenesis, we investigated protein-protein interactions within these 86 cross-disorder piDNM-containing genes. Our results revealed that 56 out of 86 (65.12%) cross-disorder genes represent an interconnected network on the level of direct/indirect protein-protein interaction relationships (Figure 5A; Supplementary Table 11). Furthermore, we determined several crucial hub piDNM-containing genes in the protein-protein interaction network, such as NOTCH1, MTOR, RYR2, and GNAS (Figure 5A), which may control common biological processes among these four neuropsychiatric disorders. Besides, these 86 cross-disorder proteins are indeed enriched in nervous system phenotypes, including abnormal synaptic transmission, abnormal nervous system development, abnormal neuron morphology and abnormal brain morphology, and behavior/neurological phenotype such as abnormal motor coordination/balance (*P* <0.05, Supplementary Figure 9A). Similarly, these 56 genes in interaction network are enriched in nervous system phenotype including abnormal nervous system development and abnormal brain morphology (*P* <0.05, Supplementary Figure 9B). By investigating the expression of these interacting genes in the human cortex of 12 ASD patients and 13 normal donators from public datasets of GSE64018^41^ and GSE76852^42^, we identified 45 (80.35%) of these PPI genes were significantly differentially expressed between ASD patients and normal controls (Student’s t-test, q <0.05, Figure 5B). And 40 (71.42%) of these PPI genes were up-regulated in ASD patients while three (5.35%) genes were down-regulated in ASD patients when compared with normal controls (Supplementary Table 12), indicating that the majority of these PPI genes were abnormally expressed in ASD patients compared with normal controls.

**Figure 5.**
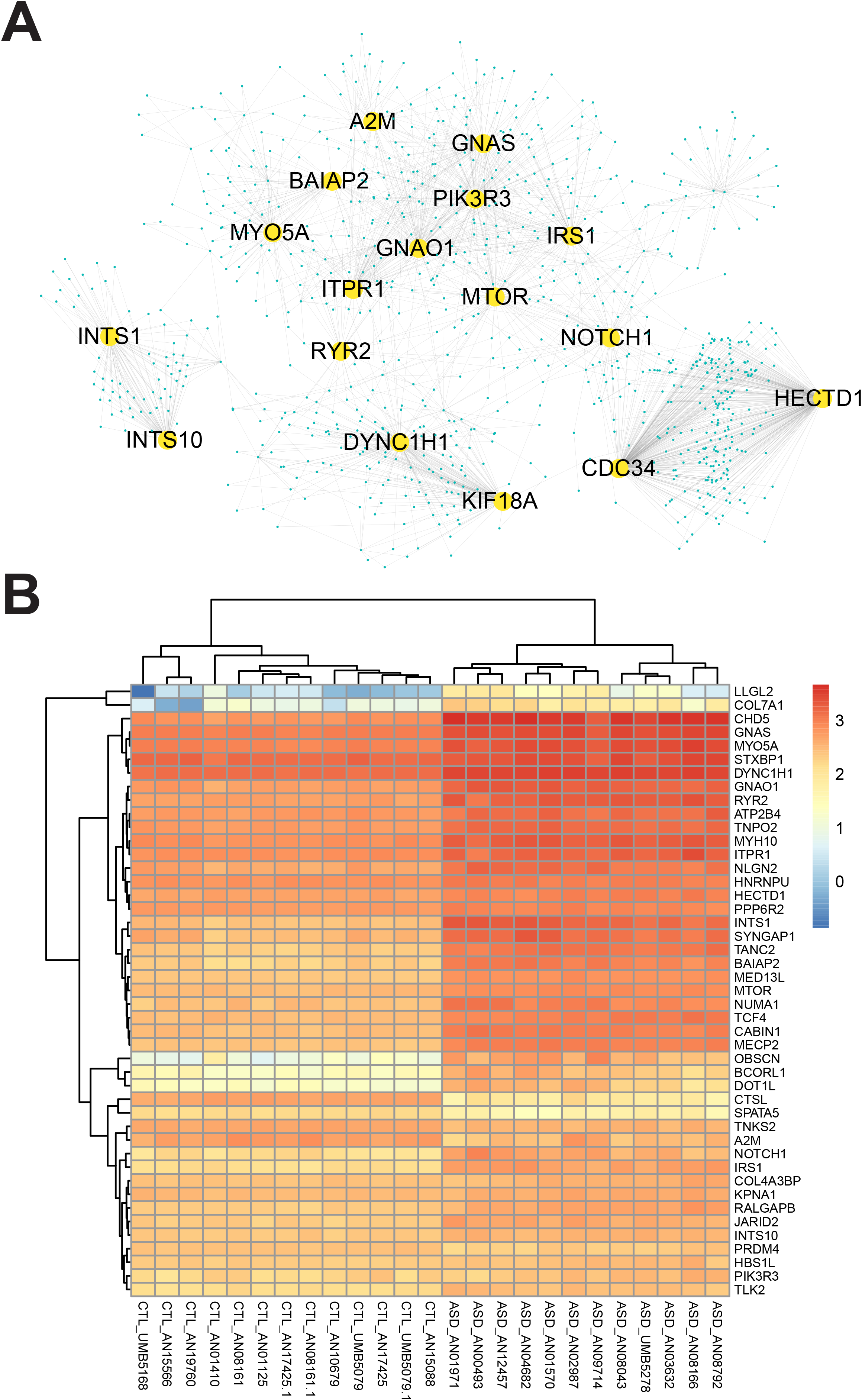
Protein-protein interaction network of the cross-disorder genes. The network of interactions between pairs of the proteins encoded by the 56 out of 86 cross-disorder genes.

### Regulatory networks between RBPs and targeting genes are potentially disrupted by piDNMs

Dysregulation or mutations of RBPs can cause a range of developmental and neurological diseases^43, 44^. Meanwhile, mutations in RNA targets of RBPs, which could disturb the interactions between RBPs and their mRNA targets, and affect mRNA metabolism and protein homeostasis in neurons during the progression of neuropathological disorders^45–47^. Hence, we constructed a regulatory network between piDNMs and RBPs based on predicted binding sites of RBPs to investigate the genetic perturbations of mRNA-RBP interactions in the four disorders (Figure 6). We identified several crucial RBP hubs that may contribute to the pathogenesis of the four neuropsychiatric disorders, including EIF4A3, FMR1, PTBP1, AGO1/2, ELAVL1, IGF2BP1/3 and WDR33. Genes with piDNMs in different disorders could be regulated by the same RBP hub while one candidate gene may be regulated by different RBP hubs (Figure 6). In addition, all of these RBP hubs were highly expressed in early fetal development stages (8-37 postconceptional weeks) based on BrainSpan developmental transcriptome (Supplementary Figure 10), suggesting their essential roles in the early stages of brain development.

**Figure 6.**
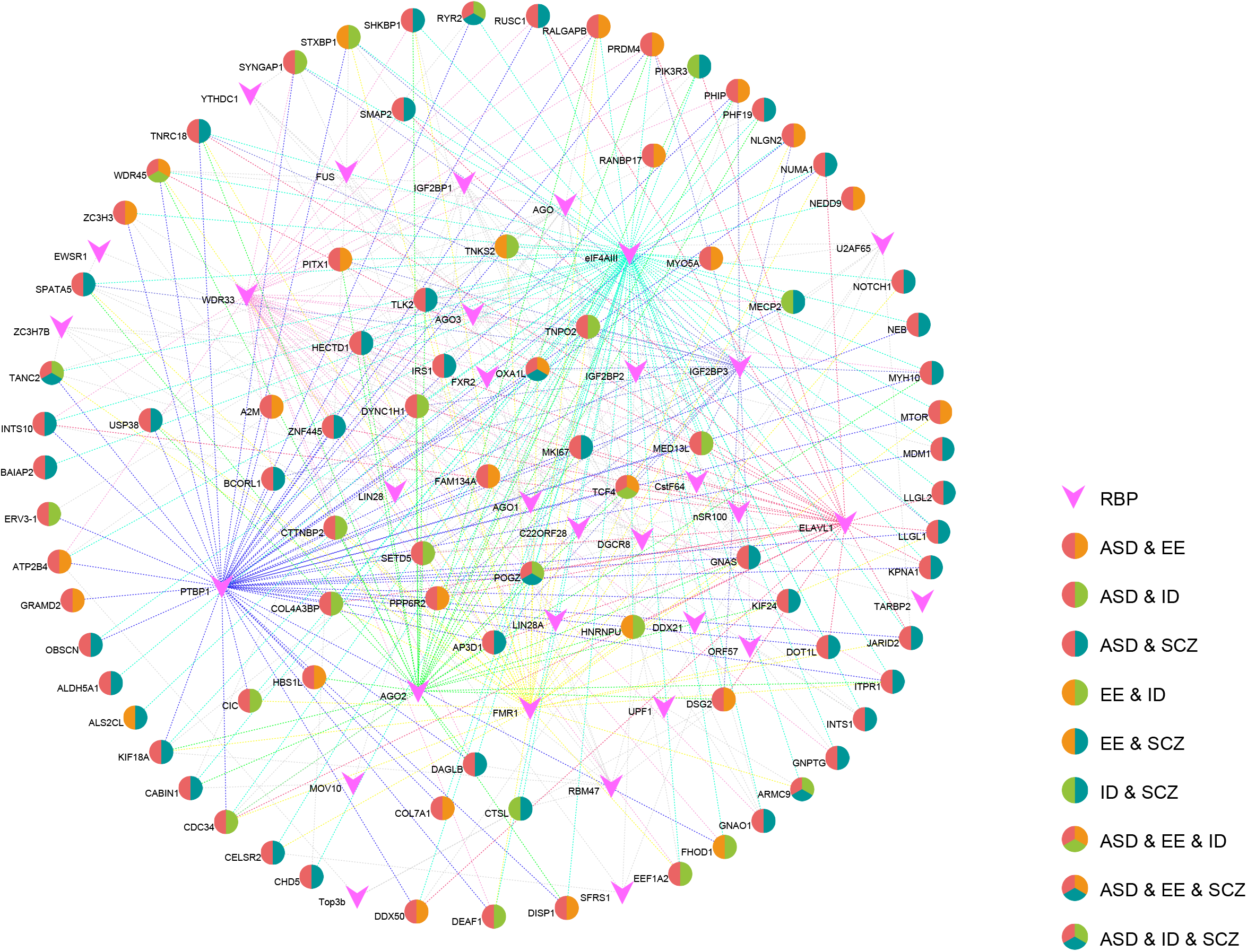
Interaction network of RBPs and genes with piDNMs. Different roles of the nodes are reflected by distinguishable geometric shapes and colors. The magenta vertical arrow stands for the RNA binding proteins. Disks with different colors represent the genes with piDNMs involved in different kinds of disorders.

## Discussion

In contrasting with the recognized role of LoF DNMs in conferring risk for neuropsychiatric disorders, the effect of DNMs on post-transcriptional regulation in pathogenesis of these disorders remains unknown. In this study, we systematically analyzed the damaging effect of DNMs on post-transcriptional regulation in four neuropsychiatric disorders, and observed higher prevalence of piDNMs in probands than that in controls in four kinds of neuropsychiatric disorders.

To date, it has been a challenge to estimate the functions of synonymous and UTRs mutations though such mutations have been widely acknowledged to alter protein expression, conformation and function^48^. We applied RBP-Var2 algorithm to annotate and interpret *de novo* variants in subjects with four neuropsychiatric disorders based on their impact to RNA secondary structure, the binding of miRNAs and RBPs. In comparison with accuracy of other prediction algorithms such as SIFT, PPH2 or RegulomeDB, RBP-Var2 has highest accuracy (AUC: 82.89%) to differentiate affected from the control subjects. Our RBP-Var2 tool identified 399 synonymous DNMs and 25 UTR’s DNMs, which were extremely harmful in post-transcriptional regulation. Consistent with previous study^49^, synonymous damaging DNMs were significantly prominent in probands compared with that in controls. Meanwhile, *de novo* insertions and deletions (InDels), especially frameshift patterns are taken for granted to be deleterious. Indeed, *de novo* frameshift InDels are more frequent in neuropsychiatric disorders compared to non-frameshift InDels^50^, which were demonstrated by predictions of RBP-Var2 but not SIFT or PPH2. Therefore, the updated version of RBP-Var2 will held great promise for exploring the effect of mutations on post-transcriptional regulation, and deciphering multiple biological layers of deleteriousness may improve the accuracy to predict disease related genetic variations.

Most interestingly, we found that some epigenetic pathways are enriched among these piDNM-containing genes, such as those that regulation of gene expression and histone modification. This finding is consistent with a previous report in which more than 68% of ASD cases shared a common acetylome aberrations at >5,000 cis-regulatory regions in prefrontal and temporal cortex^51^. Such common “epimutations” may be induced by either perturbations of epigenetic regulations, including post-transcriptional regulations due to mutations of substrates or the disruptions of epigenetic modifications resulting from the mutation of epigenetic genes. Our observations revealed the association of alterations of “epimutations” with dysregulation of post-transcription. This hypothesis is consistent with the observation that several recurrent piDNM-containing genes are non-epigenetic genes, including SYNGAP1, ADNP, POGZ and ANK2. Moreover, we discovered several recurrent epigenetic genes which contain piDNMs, including CHD8, EP300, KMT2A, KMT2C, KDM3B, JARID2 and MECP2, and they may play important roles in the genome-wide aberrations of epigenetic landscapes through disruption of the post-transcriptional regulation. Furthermore, WGCNA analysis revealed that major hubs of the co-expression network for these 86 piDNM-containing genes were histone modifiers by using BrainSpan developmental transcriptome. These data indicate that piDNM-containing genes are co-expressed with genes frequently involved in epigenetic regulation of common cellular and biological process in neuropsychiatric disorders. Importantly, these 86 piDNM-containing genes harbor intensive protein-protein interactions in physics, and shared regulatory networks between piDNMs and RBPs in four neuropsychiatric disorders. We identified several RBP hubs of regulatory networks between piDNM-containing genes and RBP proteins, including EIF4A3, FMRP, PTBP1, AGO1/2, ELAVL1, IGF2BP1/3, WDR33 and FXR2. Taking FMRP for example, it is a well-known pathogenic gene of Fragile X syndrome which co-occurs with autism in many cases and its targets are highly enriched for DNMs in ASD^52^. Our results demonstrated that, like the mutations on RBP hubs, mutations of RBP-targeting genes through disrupting their interactions with multiple RBPs may synergistically result in pathogenesis of multiple neuropsychiatric disorders.

Alterations in expression or mutations in either RBPs or their binding sites in target transcripts have been reported to cause several human diseases such as muscular atrophies, neurological disorders and cancer^53^. Although we identified 1,736 piDNMs associated with neuropsychiatric disorders, the cause and explicit effects of these piDNMs in these disorders need to be further validated and explored. In this study, our method sheds light on evaluation of post-transcriptional impact of genetic mutations especially for synonymous mutations. Additionally, as small molecules can be rapidly designed to selectively target RNAs and affect RNA-RBP interactions^54^, our study provides new insights into RNA-based therapeutic strategies for the treatment of neuropsychiatric disorders.

## Materials and methods

### Data collection and filtration

For this study, 7,748 trios or quartets were recruited from previous whole exome sequencing (WES) studies (ref), comprising 5,677 parent-probands trios associated with four neuropsychiatric disorders and 2,071 control trios (Supplementary Table 1). After removing the overlap of DNMs between probands and controls, a total of 6,996 DNMs in probands and 2,523 DNMs in controls were identified for subsequent analysis.

### RBP-Var2 algorithm

To better interpret the catalog of DNMs, we developed a new heuristic scoring system according to the functional confidence of variants based on experimental data of GWAS, eQTL, CLIP-seq derived RBP binding sites, RNA editing and miRNA targets, and machine learning algorithms. The scoring system represents with increasing confidence if a variant lies in more functional elements^14^. For example, we consider variants that are known eQTLs as significant and label them as category 1. Within category 1, subcategories indicate additional annotations ranging from the most informational variants (1a, variant may change the motif for RBP binding) to the least informational variants (1e, variant only has a motif for RBP binding). In mathematical algorithms, we employed LS-GKM^55^ (10-mer) and deltaSVM^56^ to predict the impact of DNMs on the binding of specific RBPs by calculating the delta SVM scores. Moreover, for single-base mutations, we employed the RNAsnp^57^ with default parameters to estimate the mutation effects on local RNA secondary structure and calculated the empirical *P* values based on the base pair probabilities of the wild-type and mutant RNA sequences. For insertions and deletions, we evaluated the effects of DNMs on RNA secondary structure using the minimal free energy generated by RNAfold^58^ to calculate empirical *P* values based on cumulative probabilities of the Poisson distribution. Only the functional DNM produces >5 change in gkm-SVM scores for the effect of RBP binding, and *P*-value < 0.1 or free energy change >1 for the effect of DNMs on RNA secondary structure change were determined to be a piDNM. Only DNMs occurred in exonic or UTR regions were included in our analysis.

### Identification of piDNMs and comparison with variants predicted by other methods

To determine the likelihood of a functional mutation in post-transcriptional regulation for all SNVs and InDels, our newly updated program RBP-Var2 was utilized to assign an exclusive rank for each mutation and only those mutations categorized into rank 1 or 2 were considered as piDNMs. In comparison with those mutations involved in the disruption of gene function or transcriptional regulation, several programs such as SIFT, PolyPhen2 and RegulomeDB were used to analyze the same dataset of DNMs as the input for RBP-Var2. We only kept the mutations qualified as “damaging” from the result of SIFT and “possibly damaging” or “probably damaging” from PolyPhen2. In the case of RegulomeDB, mutations labeled as category 1 and 2 were retained. Next, we classified the type of mutation (frameshift, nonframeshift, nonsynonymous, synonymous, splicing and stop) and located regions (UTR3, UTR5, exonic, ncRNA exonic and splicing) to determine the distribution of piDNMs, genetic variants and other regulatory variants. The number of variants in cases versus controls was illustrated by bar chart (***: *P*< 0.001, **: 0.001 <*P*< 0.01, *: 0.01 <*P*< 0.05, binomial test).

### TADA analysis of DNMs in four disorders

Our previously published TADA program^29^, which predicts risk genes accurately on the basis of allele frequencies, gene-specific penetrance, and mutation rate, was used to calculate the *P* value for the likelihood of each gene contributing to the all four disorders with default parameters.

### ROC curves and specificity/sensitivity estimation

We screened a positive (non-neutral) test set of likely casual mutations in Mendelian disease from the ClinVar database (v20170130). From a total of 237,308 mutations in ClinVar database, we picked up 145 exonic mutations presented in our curated DNMs in probands. Our negative (neutral) set of likely non-casual variants was built from DNMs of unaffected controls in four neuropsychiatric disorders. To exclude rare deleterious DNMs, we selected only DNMs in controls with a minor allele frequency of at least 0.01 in 1000 genome (1000g2014oct), and obtained a set of 921 exonic variants. Then, we employed R package pROC to analyze and compare ROC curves.

### Permutation analysis for overlaps of genes with piDNMs

In order to evaluate the overlap of genes among any two set of genes with piDNMs, we shuffled the intersections of genes and repeated this procedure 100,000 times. During each permutation, we randomly selected the same number of genes as the actual situation from the all RefSeq genes for each disorder taking account of gene-level de novo mutation rate, then *P* values were calculated as the proportion of permutations during which the simulated number of overlap was greater than or equal to the actual observed number.

### Functional enrichment analysis

A gene harboring piDNMs was selected into our candidate gene set to conduct functional enrichment analysis if it occurred in at least two of the four disorders. GO (Gene Ontology) and KEGG (Kyoto Encyclopedia of Genes and Genomes) pathway enrichments analyses were implemented by Cytoscape (version 3.4.0) plugin ClueGO (version 2.3.0) using genome-wide coding genes as background and *P* values calculated by hypergeometric test were corrected to be q values by Benjamini–Hochberg procedure for reducing the false discovery rate resulted from multiple hypothesis testing.

### Co-expression and spatiotemporal specificity

Normalized gene-expression of 16 human brain regions were determined by RNA sequencing and obtained from database BrainSpan (http://www.brainspan.org). We extracted expression for 77 out of 86 extreme damaging cross-disorder genes and employed R-package WGCNA (weighted correlation network analysis) with a power of five to cluster the spatiotemporal-expression patterns and prenatal laminar-expression profiles. The expression level for each gene and development stage (only stages with expression data for all 16 structures were selected, n = 14) was presented across all brain regions.

### Protein-protein interaction network of cross-disorder genes

Protein-protein interactions data of Homo sapiens was collected from the STRING (v10.5) database with score over 0.8. For the PPI network of all cross-disorder genes, we only retain the proteins with at least two links. Those nodes with degree over 30 in the network were considered as hubs. Cytoscape (version3.4.0) was used to analyze and visualize protein-protein interaction networks. Overrepresentation of mouse-mutant phenotypes was evaluated by the web tool MamPhea for the genes in the PPI network and for all cross-disorder genes containing piDNMs. The rest of genome was used as background and multiple test adjustment for *P* values was done by Benjamini-Hochberg method.

### Gene-RBP interaction network

Cytoscape (version 3.4.0) was utilized for visualization of the associations between genes harboring piDNMs in the four neuropsychiatric disorders and the corresponding regulatory RBPs.

### The available data resources

To make our findings easily accessible to the research community, we have developed RBP-Var2 platform (http://www.rbp-var.biols.ac.cn/) for storage and retrieval of piDNMs, candidate genes, and for exploring the genetic etiology of neuropsychiatric disorders in post-transcriptional regulation. The expression and epigenetic profiles of genes related to regulatory *de novo* mutations and early embryonic development have been deposited in our previously published database EpiDenovo (http://www.epidenovo.biols.ac.cn/)^40^.

## URLs

RBP-Var2, http://www.rbp-var.biols.ac.cn/; NPdenovo, http://www.wzgenomics.cn/NPdenovo/index.php; EpiDenovo: http://www.epidenovo.biols.ac.cn/; BioGRID, https://thebiogrid.org/;

MamPhea, http://evol.nhri.org.tw/phenome/index.jsp?platform; BrainSpan, http://www.brainspan.org; ClinVar, https://www.ncbi.nlm.nih.gov/clinvar/; 1000Genomes, http://www.internationalgenome.org/; WGCNA, https://labs.genetics.ucla.edu/horvath/CoexpressionNetwork/Rpackages/WGCNA/; esyN, http://www.esyn.org/; Cytoscape, http://www.cytoscape.org/; TADA, http://wpicr.wpic.pitt.edu/WPICCompGen/TADA/TADA_homepage.htm; ClueGO, http://apps.cytoscape.org/apps/cluego; pROC, http://web.expasy.org/pROC/; R, https://www.r-project.org/; Perl, https://www.perl.org/.

## Contributions

F.M. and L.W. participated in the design and execution of analyses, produced the figures, participated in the interpretation of results and edited the manuscript. F.M. developed computational code employed in the analyses. L.W. and Z.L. developed the statistical framework and drew the figures. X.Z. participated in the interpretation of results, the oversight of analyses. L.X. developed and improved the online platform of RBP-Var2. H.T. and R.C.R. provided professional guidance in the writing and refining of the manuscript. J.L. and H. T. collected the DNMs from literature and database. Z.S.S. and X.H. conceived the study, participated in the design of analyses, oversaw the study and the interpretation of results, and drafted and edited the manuscript.

## Acknowledgments

The project was funded by National Key R&D Program of China (No. 2016YFC0900400). We thank Kun Zhang for his help in TADA analysis and thank Leisheng Shi for his help in data analysis.

---

**Supplementary Figure 1**. Excess of piDNMs in probands. The odds ratio of synonymous DNMs and piDNMs were analyzed. The dominance of filtered piDNMs that not contained LoF mutations were also displayed.

**Supplementary Figure 2**. ROC curve showing the performance of the predictions of SIFT, PPH2, RBP-Var2 and RegulomeDB.

**Supplementary Figure 3**. Overlap of DNMs identified by different tools. (A) Venn diagram depicting the overlap between the DNMs predicted by SIFT, PPH2, RBP-Var2 and RegulomeDB. (B) Venn diagram depicting the overlap between the genes predicted by SIFT, PPH2, RBP-Var2 and RegulomeDB. (C) The pie chart shows the distribution of all non-LoF piDNMs. The non-LoF piDNMs detected by RBP-Var2 alone account for 52.8% of all non-LoF piDNMs (pink), while the non-LoF deleterious DNMs identified by both SIFT and Polyphen2 take up 26.2% of all (light purple). (D) Pathway enrichment analysis of the 665 genes unique to the prediction of RBP-Var2.

**Supplementary Figure 4**. Permutation test of the randomness of the overlap of different set of disease genes with control. (A-K) Permutation test for the validity of the gene overlap between the cross-disorder genes and the control. (L-O) Permutation for the overlap of genes from each disorder with control. We shuffled the genes of each disorder and calculated the shared genes between each pair, and repeated this procedure for 100,000 times to get the null distribution. The vertical dash line stands for the observed value.

**Supplementary Figure 5**. Test of the significance of the number of cross-disorder genes involved in the four neuropsychiatric disorders. (A-J) Permutation test for the validity of the gene overlap among/between every combination of three/two disorders.

**Supplementary Figure 6**. Pie chart of the pathway enrichment analysis for the 86 cross-disorder genes.

**Supplementary Figure 7**. Interaction network of the gene enrichment analysis for the 86 cross-disorder genes.

**Supplementary Figure 8**. Relationship between Co-expression modules. (A) MDS plot of genes in turquoise module and blue module. (B) Relationship between module eigengenes. (C) Clustering tree based of the module eigengenes. (D) heatmap of adjacency Eigengene.

**Supplementary Figure 9**. Mammalian phenotype enrichment analysis of selected genes. (A) Mammalian phenotype enrichment of 86 cross-disorder piDNMs genes. (B) Mammalian phenotype enrichment of 56 genes in interaction network.

**Supplementary Figure 10**. Heat map of the expression of the crucial RBP hub genes during the early fetal development stages.

## Reference

1. Iossifov I, O’Roak BJ, Sanders SJ, Ronemus M, Krumm N, Levy D et al. The contribution of de novo coding mutations to autism spectrum disorder. Nature 2014; 515(7526): 216–221.

2. Pires DE, Ascher DB, Blundell TL. DUET: a server for predicting effects of mutations on protein stability using an integrated computational approach. Nucleic Acids Res 2014; 42(Web Server issue): W314–319.

3. Adzhubei IA, Schmidt S, Peshkin L, Ramensky VE, Gerasimova A, Bork P et al. A method and server for predicting damaging missense mutations. Nature Methods 2010; 7(4): 248–249.

4. Chen XF, Zhang YW, Xu HX, Bu GJ. Transcriptional regulation and its misregulation in Alzheimer’s disease. Molecular Brain 2013; 6.

5. Lee TI, Young RA. Transcriptional Regulation and Its Misregulation in Disease. Cell 2013; 152(6): 1237–1251.

6. Portela A, Esteller M. Epigenetic modifications and human disease. Nature Biotechnology 2010; 28(10): 1057–1068.

7. Sakabe NJ, Savic D, Nobrega MA. Transcriptional enhancers in development and disease. Genome Biology 2012; 13(1).

8. Boyle AP, Hong EL, Hariharan M, Cheng Y, Schaub MA, Kasowski M et al. Annotation of functional variation in personal genomes using RegulomeDB. Genome Res 2012; 22(9): 1790–1797.

9. Wan Y, Qu K, Zhang QC, Flynn RA, Manor O, Ouyang Z et al. Landscape and variation of RNA secondary structure across the human transcriptome. Nature 2014; 505(7485): 706–709.

10. Suhl JA, Muddashetty RS, Anderson BR, Ifrim MF, Visootsak J, Bassell GJ et al. A 3’ untranslated region variant in FMR1 eliminates neuronal activity-dependent translation of FMRP by disrupting binding of the RNA-binding protein HuR. Proc Natl Acad Sci U S A 2015; 112(47): E6553–6561.

11. Chin LJ, Ratner E, Leng S, Zhai R, Nallur S, Babar I et al. A SNP in a let-7 microRNA complementary site in the KRAS 3’ untranslated region increases non-small cell lung cancer risk. Cancer Res 2008; 68(20): 8535–8540.

12. Maticzka D, Lange SJ, Costa F, Backofen R. GraphProt: modeling binding preferences of RNA-binding proteins. Genome Biol 2014; 15(1): R17.

13. Fukunaga T, Ozaki H, Terai G, Asai K, Iwasaki W, Kiryu H. CapR: revealing structural specificities of RNA-binding protein target recognition using CLIP-seq data. Genome Biol 2014; 15(1): R16.

14. Mao FB, Xiao LY, Li XF, Liang JL, Teng HJ, Cai WS et al. RBP-Var: a database of functional variants involved in regulation mediated by RNA-binding proteins. Nucleic Acids Research 2016; 44(D1): D154–D163.

15. Hu B, Yang Y-CT, Huang Y, Zhu Y, Lu ZJ. POSTAR: a platform for exploring post-transcriptional regulation coordinated by RNA-binding proteins. Nucleic Acids Research 2016: gkw888.

16. Elsabbagh M, Divan G, Koh YJ, Kim YS, Kauchali S, Marcin C et al. Global Prevalence of Autism and Other Pervasive Developmental Disorders. Autism Research 2012; 5(3): 160–179.

17. Gamazon ER, Badner JA, Cheng L, Zhang C, Zhang D, Cox NJ et al. Enrichment of cis-regulatory gene expression SNPs and methylation quantitative trait loci among bipolar disorder susceptibility variants. Molecular Psychiatry 2013; 18(3): 340–346.

18. Bonder MJ, Luijk R, Zhernakova DV, Moed M, Deelen P, Vermaat M et al. Disease variants alter transcription factor levels and methylation of their binding sites. Nat Genet 2016.

19. Loke YJ, Hannan AJ, Craig JM. The Role of Epigenetic Change in Autism Spectrum Disorders. Front Neurol 2015; 6: 107.

20. Gupta S, Ellis SE, Ashar FN, Moes A, Bader JS, Zhan J et al. Transcriptome analysis reveals dysregulation of innate immune response genes and neuronal activity-dependent genes in autism. Nat Commun 2014; 5: 5748.

21. Voineagu I, Wang X, Johnston P, Lowe JK, Tian Y, Horvath S et al. Transcriptomic analysis of autistic brain reveals convergent molecular pathology. Nature 2011; 474(7351): 380–384.

22. Corbett BA, Kantor AB, Schulman H, Walker WL, Lit L, Ashwood P et al. A proteomic study of serum from children with autism showing differential expression of apolipoproteins and complement proteins. Mol Psychiatry 2007; 12(3): 292–306.

23. Short PJ, McRae JF, Gallone G, Sifrim A, Won H, Geschwind DH et al. De novo mutations in regulatory elements in neurodevelopmental disorders. Nature 2018.

24. Liu Y, Liang Y, Cicek AE, Li Z, Li J, Muhle RA et al. A Statistical Framework for Mapping Risk Genes from De Novo Mutations in Whole-Genome-Sequencing Studies. The American Journal of Human Genetics 2018.

25. Zhou J, Park CY, Theesfeld CL, Wong AK, Yuan Y, Scheckel C et al. Whole-genome deep-learning analysis identifies contribution of noncoding mutations to autism risk. Nat Genet 2019; 51(6): 973–980.

26. Boyle AP, Hong EL, Hariharan M, Cheng Y, Schaub MA, Kasowski M et al. Annotation of functional variation in personal genomes using RegulomeDB. Genome Res 2012; 22(9): 1790–1797.

27. Wang W, Bu B, Xie M, Zhang M, Yu Z, Tao D. Neural cell cycle dysregulation and central nervous system diseases. Prog Neurobiol 2009; 89(1): 1–17.

28. Lau CG, Zukin RS. NMDA receptor trafficking in synaptic plasticity and neuropsychiatric disorders. Nat Rev Neurosci 2007; 8(6): 413–426.

29. He X, Sanders SJ, Liu L, De Rubeis S, Lim ET, Sutcliffe JS et al. Integrated Model of De Novo and Inherited Genetic Variants Yields Greater Power to Identify Risk Genes. Plos Genetics 2013; 9(8).

30. Li JC, Cai T, Jiang Y, Chen HQ, He X, Chen C et al. Genes with de novo mutations are shared by four neuropsychiatric disorders discovered from NPdenovo database. Molecular Psychiatry 2016; 21(2): 290–297.

31. Brookes E, Shi Y. Diverse Epigenetic Mechanisms of Human Disease. Annual Review of Genetics, Vol 48 2014; 48: 237–268.

32. Peter CJ, Akbarian S. Balancing histone methylation activities in psychiatric disorders. Trends in Molecular Medicine 2011; 17(7): 372–379.

33. Egan CM, Nyman U, Skotte J, Streubel G, Turner S, O’Connell DJ et al. CHD5 Is Required for Neurogenesis and Has a Dual Role in Facilitating Gene Expression and Polycomb Gene Repression. Developmental Cell 2013; 26(3): 223–236.

34. Dubey N, Hoffman JF, Schuebel K, Yuan Q, Martinez PE, Nieman LK et al. The ESC/E(Z) complex, an effector of response to ovarian steroids, manifests an intrinsic difference in cells from women with premenstrual dysphoric disorder. Mol Psychiatry 2017; 22(8): 1172–1184.

35. Chittka A, Nitarska J, Grazini U, Richardson WD. Transcription factor positive regulatory domain 4 (PRDM4) recruits protein arginine methyltransferase 5 (PRMT5) to mediate histone arginine methylation and control neural stem cell proliferation and differentiation. J Biol Chem 2012; 287(51): 42995–43006.

36. Wang H, Westin L, Nong Y, Birnbaum S, Bendor J, Brismar H et al. Norbin Is an Endogenous Regulator of Metabotropic Glutamate Receptor 5 Signaling. Science 2009; 326(5959): 1554–1557.

37. Lotan A, Fenckova M, Bralten J, Alttoa A, Dixson L, Williams RW et al. Neuroinformatic analyses of common and distinct genetic components associated with major neuropsychiatric disorders. Frontiers in Neuroscience 2014; 8.

38. Langfelder P, Horvath S. WGCNA: an R package for weighted correlation network analysis. Bmc Bioinformatics 2008; 9.

39. Kang HJ, Kawasawa YI, Cheng F, Zhu Y, Xu X, Li M et al. Spatio-temporal transcriptome of the human brain. Nature 2011; 478(7370): 483–489.

40. Mao F, Liu Q, Zhao X, Yang H, Guo S, Xiao L et al. EpiDenovo: a platform for linking regulatory de novo mutations to developmental epigenetics and diseases. Nucleic Acids Res 2017.

41. Irimia M, Weatheritt RJ, Ellis JD, Parikshak NN, Gonatopoulos-Pournatzis T, Babor M et al. A Highly Conserved Program of Neuronal Microexons Is Misregulated in Autistic Brains. Cell 2014; 159(7): 1511–1523.

42. Werling DM, Parikshak NN, Geschwind DH. Gene expression in human brain implicates sexually dimorphic pathways in autism spectrum disorders. Nature Communications 2016; 7.

43. Castello A, Fischer B, Hentze MW, Preiss T. RNA-binding proteins in Mendelian disease. Trends in Genetics 2013; 29(5): 318–327.

44. Klein ME, Monday H, Jordan BA. Proteostasis and RNA Binding Proteins in Synaptic Plasticity and in the Pathogenesis of Neuropsychiatric Disorders. Neural Plasticity 2016.

45. Lukong KE, Chang KW, Khandjian EW, Richard S. RNA-binding proteins in human genetic disease. Trends in Genetics 2008; 24(8): 416–425.

46. Gerstberger S, Hafner M, Tuschl T. A census of human RNA-binding proteins. Nature Reviews Genetics 2014; 15(12): 829–845.

47. Neelamraju Y, Hashemikhabir S, Janga SC. The human RBPome: From genes and proteins to human disease. Journal of Proteomics 2015; 127: 61–70.

48. Sauna ZE, Kimchi-Sarfaty C. Understanding the contribution of synonymous mutations to human disease. Nature Reviews Genetics 2011; 12(10): 683–691.

49. Takata A, Ionita-Laza I, Gogos JA, Xu B, Karayiorgou M. De Novo Synonymous Mutations in Regulatory Elements Contribute to the Genetic Etiology of Autism and Schizophrenia. Neuron 2016; 89(5): 940–947.

50. Kosmicki JA, Samocha KE, Howrigan DP, Sanders SJ, Slowikowski K, Lek M et al. Refining the role of de novo protein-truncating variants in neurodevelopmental disorders by using population reference samples. Nat Genet 2017.

51. Sun W, Poschmann J, Cruz-Herrera Del Rosario R, Parikshak NN, Hajan HS, Kumar V et al. Histone Acetylome-wide Association Study of Autism Spectrum Disorder. Cell 2016; 167(5): 1385–1397 e1311.

52. Parikshak NN, Gandal MJ, Geschwind DH. Systems biology and gene networks in neurodevelopmental and neurodegenerative disorders. Nature Reviews Genetics 2015; 16(8): 441–458.

53. Kechavarzi B, Janga SC. Dissecting the expression landscape of RNA-binding proteins in human cancers. Genome Biology 2014; 15(1).

54. Costales MG, Haga CL, Velagapudi SP, Childs-Disney JL, Phinney DG, Disney MD. Small Molecule Inhibition of microRNA-210 Reprograms an Oncogenic Hypoxic Circuit. J Am Chem Soc 2017; 139(9): 3446–3455.

55. Lee D. LS-GKM: a new gkm-SVM for large-scale datasets. Bioinformatics 2016; 32(14): 2196–2198.

56. Lee D, Gorkin DU, Baker M, Strober BJ, Asoni AL, McCallion AS et al. A method to predict the impact of regulatory variants from DNA sequence. Nature Genetics 2015; 47(8): 955-+.

57. Sabarinathan R, Tafer H, Seemann SE, Hofacker IL, Stadler PF, Gorodkin J. RNAsnp: Efficient Detection of Local RNA Secondary Structure Changes Induced by SNPs (vol 34, pg 546, 2013). Human Mutation 2013; 34(6): 925–925.

58. Gruber AR, Lorenz R, Bernhart SH, Neuboock R, Hofacker IL. The Vienna RNA Websuite. Nucleic Acids Research 2008; 36: W70–W74.

